# Highly pathogenic avian influenza (HPAI) in South America, 2022-2025: temporality, affected species, and the southwards expansion to the Antarctic region

**DOI:** 10.1101/2025.10.03.680239

**Authors:** Fernanda Sánchez-Rodríguez, Constanza Diaz-Gavidia, Soledad Ruíz, Pedro Jimenez-Bluhm

**Affiliations:** Escuela de Medicina, Facultad de Medicina, Facultad de Ciencias Biológicas y Facultad de Agronomía y Sistemas Naturales, Pontificia Universidad Católica de Chile, Santiago 7820436, Chile; Escuela de Medicina Veterinaria, Facultad de Recursos Naturales y Medicina Veterinaria, Universidad Santo Tomás, Santiago 8820000, Chile

## Abstract

The H5N1 highly pathogenic avian influenza (HPAI) virus has caused severe global losses, reaching South America in 2022 and Antarctica in 2024. Here we synthesize outbreak reports submitted to the World Organization for Animal Health (WOAH) by South American countries and document the virus’s unprecedented expansion into Antarctica, affecting wild birds, wild mammals, and domestic poultry. More than 6 million domestic birds died or were culled, mostly from commercial operations. Of the 11 South American countries that reported H5N1 to WOAH, 10 reported infections in wild birds, spanning 104 species, 59.62% of which are migratory and predominantly non-trans-equatorial. Marine mammal cases occurred after wild bird detections, with the South American sea lion (*Otaria flavescens*) most affected, and several Antarctic bird species with migratory behavior were also reported in South America. To complement outbreak data, we examined available genomic sequences through phylogenetic and time-calibrated Bayesian analyses, which revealed multiple introduction events, viral diversity across regions, and evidence of interspecies transmission dynamics. These findings highlight the extensive ecological reach of H5N1 in the Southern Hemisphere and underscore the urgent need for a One Health approach that strengthens wildlife and backyard-poultry surveillance while fostering coordinated regional action to control and prevent further spread of HPAI.

**IMPORTANCE:** The arrival of H5N1 highly pathogenic avian influenza (HPAI) in South America has caused severe mortality in wild birds, marine mammals, and domestic poultry, and has recently expanded into Antarctica. Understanding how the virus entered and spread across the continent is essential for preparedness and response. Using phylogenetic and time-calibrated analyses, we identify three independent introductions into South America, estimate their temporal windows of entry, and document repeated spillover across species, including into marine mammals and humans. These findings provide novel resolution beyond previous reports and highlight the extensive inter-country connectivity of circulating viruses. The unprecedented detection of HPAI in Antarctica further illustrates the ecological risks posed by ongoing southward spread. Together, this analysis underscores the urgent need for integrated One Health surveillance that bridges wildlife, domestic animal, and human health systems to mitigate the future impacts of HPAI in the region.

## INTRODUCTION

Influenza A virus (IAV), a member of the Orthomyxoviridae family, is a highly adaptable pathogen with a broad host range and significant potential for cross-species transmission (1, 2). It is primarily maintained in nature by aquatic wild birds, which serve as reservoirs for diverse hemagglutinin (HA) and neuraminidase (NA) subtype combinations (3). Influenza A viruses are classified based on variations in these surface proteins. To date, 17 distinct antigenic variants of the HA and 9 different NA subtypes have been identified in avian species (4). In addition to antigenic diversity, IAVs are categorized by their pathogenic phenotypes in poultry. Although many strains are of low pathogenicity (LPAI) and typically cause mild or no clinical signs, they may still contribute to disease under certain environmental conditions or in combination with secondary infections (5). In contrast, highly pathogenic avian influenza viruses (HPAI) can cause mortality rates approaching 100% in chickens, a phenotype that has so far only been observed in H5 and H7 subtypes (6). Genetic mutations or reassortment events can transform LPAI into HPAI, resulting in severe disease in poultry, wild birds, and occasionally mammals (7). A key molecular marker of this transition is the acquisition of a polybasic cleavage site in the precursor HA protein (HA0), which permits cleavage by ubiquitous host proteases rather than being limited to trypsin-like enzymes in specific anatomical regions. This enables HPAI to replicate systemically, contributing to its heightened virulence (8). The polybasic cleavage site can arise through mechanisms such as gene insertion, substitution, or recombination with other viral gene segments (6, 9, 10). Beyond point mutations, the segmented genome of IAV enables genetic reassortment during coinfections, producing novel viruses without pre-existing population immunity (11). Its adaptability, transmissibility, and pathogenicity arise from viral– host interactions, with the low-fidelity RNA-dependent RNA polymerase generating frequent mutations that, under selective pressures, can enhance transmissibility, virulence, or drug resistance (12, 13). These molecular features underpin the virus’s capacity for long-distance spread and host switching, as illustrated by the global emergence and diversification of HPAI H5N1 clade 2.3.4.4b (14).

The emergence of HPAI H5N1 clade 2.3.4.4b, which has recently spread across multiple continents, including Asia, the Middle East, Europe, and the Americas, originated from the Gs/Gd lineage of HPAIs first detected in Guangdong in 1996, through genomic reassortment events involving LPAI viruses circulating in wild bird populations (15). The Gs/Gd viruses marked a turning point in avian influenza evolution, driving diversification into multiple hemagglutinin (HA) clades and affecting both domestic and wild bird species, including humans, and causing significant economic losses and food insecurity (16, 17). Mass mortality events in wild birds began with a 2005 outbreak at Qinghai Lake, China, involving clade 2.2 viruses that spread via migratory birds (18, 19).

Since 2009, HPAI has undergone multiple reassortment events with LPAI viruses, leading to the emergence of new H5 subtypes. Among these, clade 2.3.4.4, and particularly its subgroups 2.3.4.4a and 2.3.4.4b, gained dominance in subsequent years (15). Clade 2.3.4.4a emerged in South Korea during the winter of 2013–2014 through reassortment between H5N8 strains (20). It spread via migratory birds to Russia, Europe, and North America in late 2014 through the Beringia region, marking the first intercontinental transmission of HPAI H5N8 (21). However, the virus was eradicated from North America by mid-2016 (22).

In contrast, clade 2.3.4.4b emerged between May and June 2016 in wild birds located in China and along the Russia–Mongolia border, as a result of reassortment events involving H5N8, H4N2, and H11N9 viruses (20). It later spread to South Asia, Europe, and Africa, undergoing further reassortments with local LPAI viruses to produce new H5N6 and H5N5 subtypes (23). Continued viral circulation along migratory flyways led to the emergence of H5N1 in Northern Europe, marking the third major intercontinental epizootic event (24). This new dominant H5N1 subtype, first identified in the Netherlands in 2020, replaced H5N8 by 2021–2022. It originated from reassortment between H5N8 and various LPAI viruses, highlighting the complex evolution and adaptability of clade 2.3.4.4b (15, 24).

Migratory birds then introduced the virus to the American High Arctic in late 2021, first detected in Newfoundland, Canada, followed by a spread along the U.S. east coast and into the Americas (25), causing widespread deaths in wild birds and mammals. This pattern differed from earlier European epizootics, where swans were the most affected species during 2005– 2006 and 2016–2017, followed by ducks. (26). Over time, both the number of wild mammal species infected with H5N1 and the virus’s geographic range have expanded markedly. Before 2020, infections in wild mammals were confined to terrestrial and semi-aquatic species, with no cases reported in the Americas. Since 2020, however, the virus has extended its reach, with aquatic mammals now accounting for 25% of reported cases worldwide (27).

HPAI has affected many countries worldwide, and although South America has not been an exception, the number of outbreaks reported in this region has been comparatively lower, and only a limited number of distinct viral strains have been detected. To date, only two HPAI subtypes have been identified in South America: H7N3 in commercial broiler chicken in Chile in 2002 (28), and currently H5N1 clade 2.3.4.4b, which began circulating in South American countries since the second half of 2022 (29). The latter has expanded its geographic range and host spectrum, raising concerns about its ecological impact, potential for reassortment, and risk of zoonotic spillover (30). While much of the early global focus has centered on outbreaks in Europe, Asia, and North America, the progressive southward dissemination of H5N1 into South America, and even into the Antarctic region, represents an emerging frontier in the global epidemiology of HPAIV.

In this manuscript, we present an exhaustive review of publicly available data to characterize the impact of highly pathogenic avian influenza in South America. This includes a comprehensive search of reported outbreaks, affected species, and the geographic extent of the virus’s spread across the continent. Additionally, we analyze available viral genome sequences from the region to provide an integrated perspective on the genetic diversity and dynamics of the circulating strains. By combining epidemiological and genomic information, our goal is to offer a resource for stakeholders, including public health officials, wildlife conservation managers, and policymakers, to better understand the scope and drivers of these outbreaks. Such understanding is crucial for anticipating and managing future events, especially given the virus’s demonstrated capacity for rapid, transboundary, and lethal spread.

## RESULTS

### Outbreaks

Eleven out of the 14 countries in South America reported HPAI H5N1 to the WOAH either in wild birds, commercial domestic birds, backyard domestic birds, or wild mammals up to May 2025 (WOAH, 2025). No cases were reported in French Guiana, Guyana, and Suriname. Among the 11 countries that reported HPAI H5N1 cases, seven (63.64%) identified wild birds as their first affected host group (Venezuela, Chile, Argentina, Uruguay, Brazil, Peru, and the Falkland Islands). Three countries (Colombia, Bolivia, and Paraguay) initially reported cases in backyard domestic birds, while two countries (Ecuador and Bolivia) reported their first detections in commercial domestic birds. Notably, Bolivia simultaneously reported outbreaks in both commercial and backyard domestic birds. Furthermore, six out of the seven countries where wild birds were the first affected hosts, backyard poultry constituted the second group in which the virus was detected; the Falkland Islands is the only country not included here. A detailed graphical timeline showing the temporal dynamics of outbreaks across the region is presented in Figure 1.

**Figure 1.**
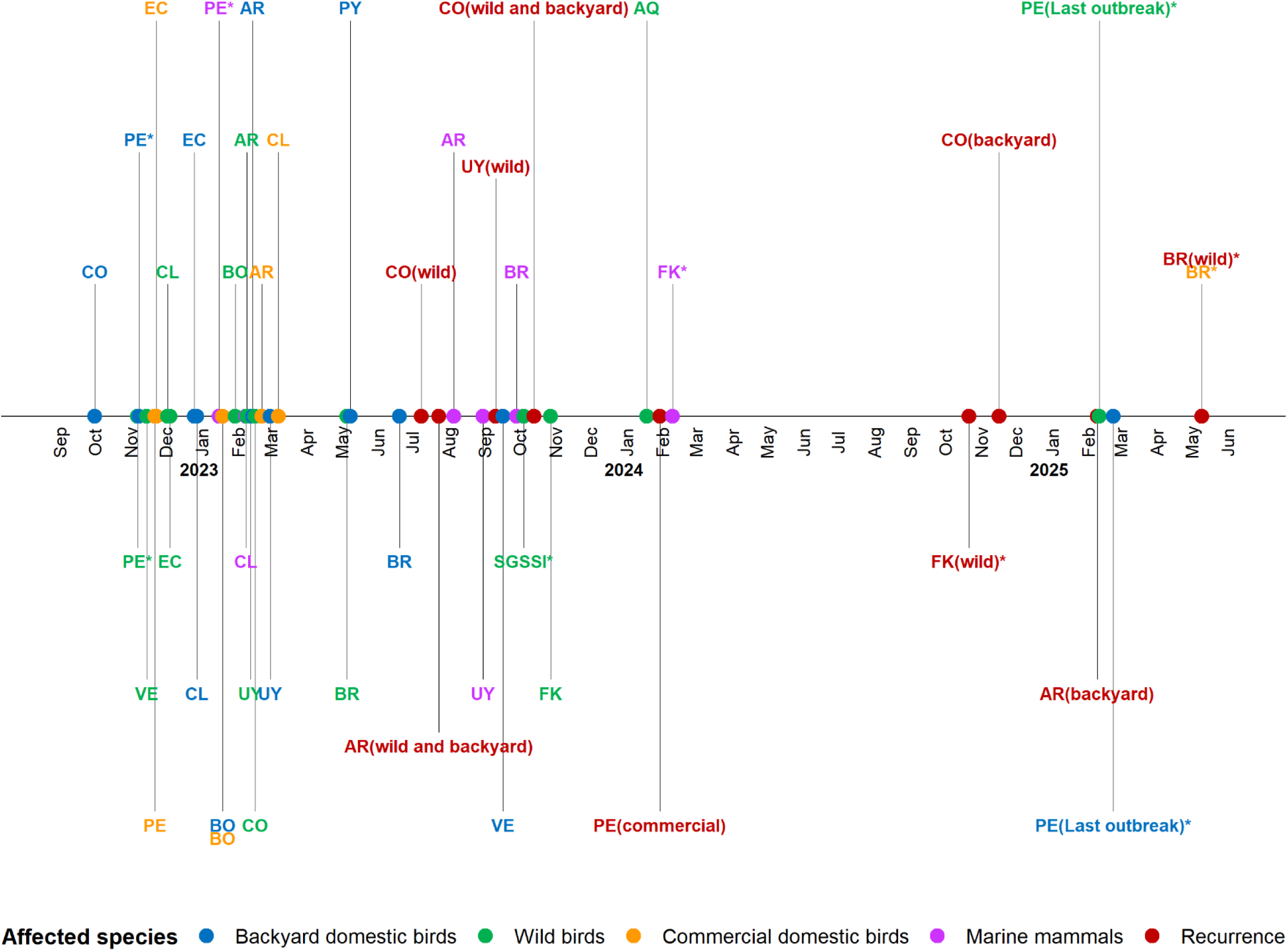
Timeline of the first highly pathogenic avian influenza cases reported to the World Organization of Animal Health (WOAH) from South American countries until May 2025, present in the World Animal Health Information System (WAHIS). All of the reported events are from the H5N1 outbreak. (AQ: Antarctica, AR: Argentina, BO: Bolivia, BR: Brazil, CO: Colombia, CL: Chile, EC: Ecuador, FK: Falkland Islands, PE: Peru, PY: Paraguay, SGSSI: South Georgia and the South Sandwich Islands, VE: Venezuela, UY: Uruguay) *These events were not closed by May 2025.

All South American countries that reported HPAI H5N1 to the WOAH documented cases in wild birds, except for Paraguay. The first wild bird species affected varied by country: brown pelicans (*Pelecanus occidentalis*) in Colombia and Venezuela; Peruvian pelicans (*Pelecanus thagus*) in Peru and Chile; blue-footed boobies (*Sula nebouxii*) in Peru and Ecuador; blue-and-white swallows (*Pygochelidon cyanoleuca*) in Bolivia; Andean geese (*Oressochen melanopterus*) in Argentina; black-necked swans (*Cygnus melancoryphus*) in Uruguay; Cabot’s terns (*Thalasseus sandvicensis* spp. *acuflavidus*) in Brazil; and southern fulmars (*Fulmarus glacialoides*) in the Falkland Islands.

A total of ten South American countries reported HPAI H5N1 cases in backyard domestic birds (Colombia, Peru, Chile, Bolivia, Ecuador, Argentina, Uruguay, Paraguay, Brazil, and Venezuela). In addition, six countries reported cases in marine mammals (Peru, Chile, Argentina, Uruguay, Brazil, and the Falkland Islands). Cases in commercial poultry were documented in six countries (Argentina, Bolivia, Brazil, Chile, Ecuador, and Peru). Paraguay exclusively reported cases in backyard domestic birds, while the Falkland Islands reported cases only in wild animals (birds and marine mammals). Notably, all marine mammal cases occurred after wild bird cases.

Several countries have exhibited recurrence of HPAI H5N1 in specific host groups. Wild birds were the most frequently affected by recurrent outbreaks, as observed in Argentina, Brazil, Colombia, Uruguay, and the Falkland Islands. Recurrence in backyard poultry was reported in Colombia and Argentina, while Peru was the only country to report recurrence in commercial poultry. The last outbreaks reported in South America, which were ongoing up to May 2025, were reported in Peru, affecting both wild birds and backyard poultry, and in Brazil, where cases persisted in wild birds. Other outbreaks that remain officially open have not reported additional cases.

### Affected species of HPAI in South America

#### Domestic birds

From 2022 to May 2025, South American countries reported a total of 2,930,410 H5N1-positive domestic birds to WOAH, with 6,147,176 deaths/eliminations recorded (Supplementary Table 1). In backyard poultry, 20,121 birds tested positive for the virus, while 51,730 died or were eliminated due to this virus. Peru reported the highest number of cases, followed by Chile, Argentina, and Colombia (Figure 2). In contrast, Chile recorded the highest number of deaths and eliminations, followed by Peru.

**Figure 2.**
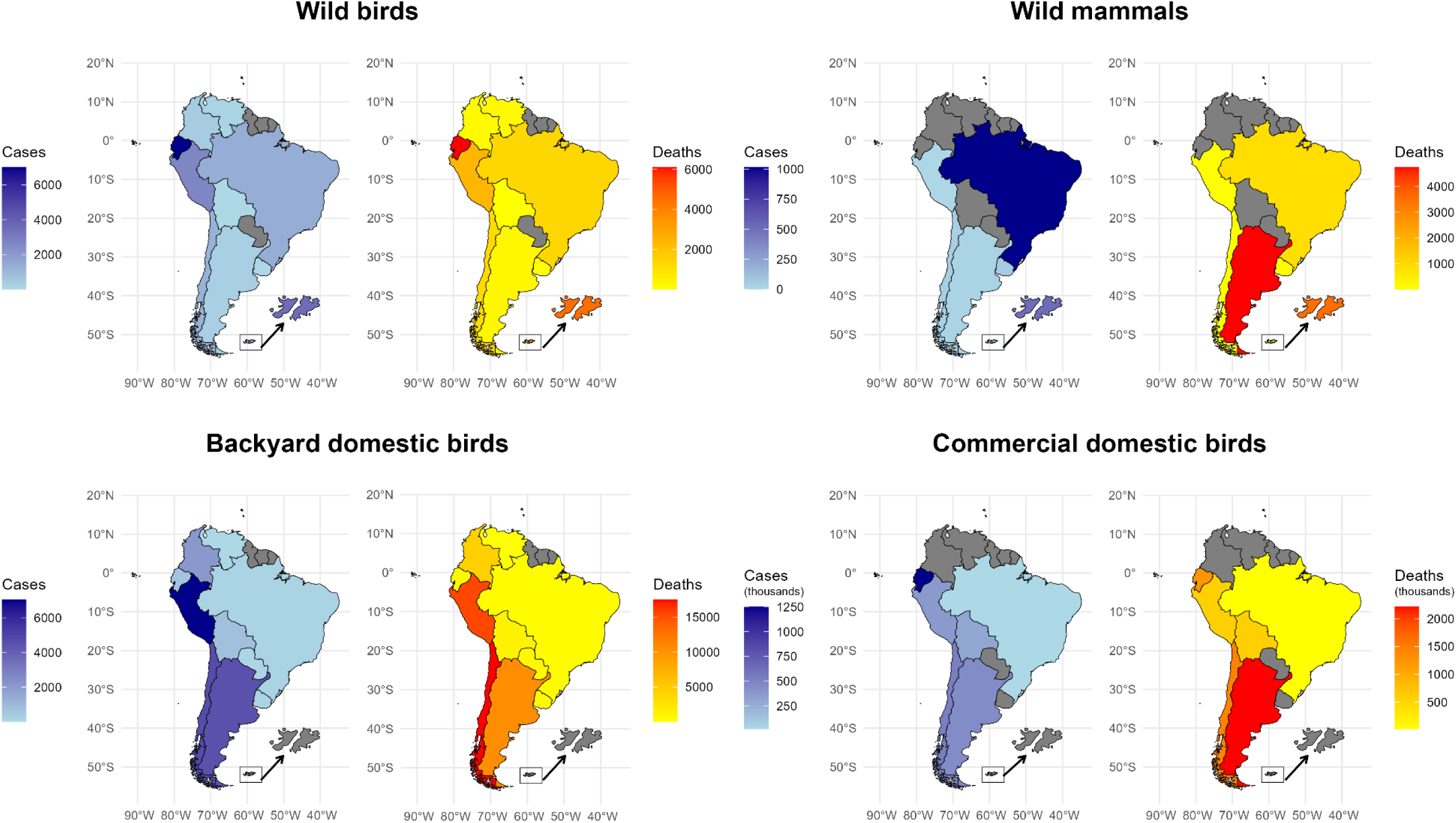
Number of positive cases, as well as deaths (including eliminations) from H5N1 reported by South American countries to the World Organization of Animal Health (WOAH), until the 30^th^ of May 2025, present in the World Animal Health Information System (WAHIS).

Regarding commercial domestic birds, more than 2.9 million birds were reported as H5N1- positive across South America, with over 6 million deaths or eliminations. Ecuador registered the highest number of positive cases, followed by Chile and Argentina (Figure 2), while Argentina reported the largest number of deaths and eliminations, followed by Chile and Ecuador.

#### Wild birds

A total of 13,118 confirmed H5N1-positive cases and 14,505 associated deaths or eliminations were recorded in wild birds across South America. In total, 104 identified wild bird species and five unidentified species were reported (Table 1). Chile reported the greatest species diversity, with 57 affected species, followed by Brazil (n = 34) and Peru (n = 27). All other countries reported fewer than 10 species.

**Table 1.** Wild bird species that were reported by South American countries to the World Organization for Animal Health (WOAH) for “Influenza A viruses of high pathogenicity” until the 30^th^ of May 2025, which are present in the World Animal Health Information System (WAHIS). The global assessment of the IUCN Red List Category was included for each species.

| Species | Scientific name | Country | Cases | Deaths | Eliminations | IUCN |
| --- | --- | --- | --- | --- | --- | --- |
| American golden plover | *Pluvialis dominica* | Brazil | 1 | 0 | 1 | Least concern |
| American oystercatcher | *Haematopus palliatus* | Chile | 1 | 1 | 0 | Least concern |
| Anatidae (unidentified) |  | Argentina | 3 | 3 | 0 |  |
| Andean condor | *Vultur gryphus* | Peru | 1 | 1 | 0 | Vulnerable |
| Andean goose | *Oressochen melanopterus* | Argentina | 2 | 2 | 0 | Least concern |
|  |  | Peru | 102 | 102 | 0 |  |
| Anhinga | *Anhinga anhinga* | Brazil | 2 | 2 | 0 | Least concern |
| Antarctic Prion | *Pachyptila desolata* | Brazil | 1 | 0 | 1 | Least concern |
| Austral parakeet | *Enicognathus ferrugineus* | Chile | 1 | 0 | 1 | Least concern |
| Belcher's gull | *Larus belcheri* | Chile | 27 | 27 | 0 | Least concern |
|  |  | Peru | 0 | 1 | 0 |  |
| Black skimmer | *Rynchops niger* | Chile | 6 | 6 | 0 | Least concern |
|  |  | Peru | 12 | 11 | 0 |  |
| Black swan | *Cygnus atratus* | Brazil | 96 | 84 | 0 | Least concern |
| Black vulture | *Coragyps atratus* | Chile | 5 | 5 | 0 | Least concern |
|  |  | Peru | 1 | 1 | 0 |  |
| Black-browed albatross | *Thalassarche melanophris* | Chile | 1 | 1 | 0 | Least concern |
|  |  | Falkland Is. | 1 | 1 | 0 |  |
| Black-chested buzzard-eagle | *Geranoaetus melanoleucus* | Chile | 1 | 1 | 0 | Least concern |
|  |  | Peru | 13 | 10 | 2 |  |
| **Black-crowned Night-Heron** | *Nycticorax nycticorax* | Chile | 2 | 2 | 0 | Least concern |
| **Black-faced ibis** | *Theristicus melanopis* | Chile | 1 | 1 | 0 | Least concern |
| **Blackish oystercatcher** | *Haematopus ater* | Chile | 4 | 4 | 0 | Least concern |
| **Black-necked swan** | *Cygnus melancoryphus* | Argentina | 21 | 19 | 0 | Least concern |
|  |  | Brazil | 141 | 132 | 9 |  |
|  |  | Chile | 123 | 119 | 4 |  |
|  |  | Uruguay | 14 | 14 | 0 |  |
| **Blue-and-white swallow** | *Pygochelidon cyanoleuca* | Bolivia | 4 | 4 | 0 | Least concern |
| **Blue-footed booby** | *Sula nebouxii* | Ecuador | 4 | 3 | 1 | Least concern |
|  |  | Peru | 4 | 2 | 1 |  |
| **Brown Booby** | *Sula leucogaster* | Brazil | 3 | 0 | 3 | Least concern |
| **Brown pelican** | *Pelecanus occidentalis* | Colombia | 376 | 167 | 1 | Least concern |
|  |  | Venezuela | 172 | 172 | 0 |  |
| **Brown skua** | *Stercorarius antarcticus* | Falkland Is. | 0 | 7 | 0 | Least concern |
| **Brown-hooded gull** | *Chroicocephalus maculipennis* | Brazil | 1 | 0 | 1 | Least concern |
|  |  | Chile | 7 | 7 | 0 |  |
|  |  | Peru | 10 | 8 | 0 |  |
| **Burrowing parrot** | *Cyanoliseus patagonus* | Chile | 2 | 0 | 2 | Least concern |
| **Cabot's tern** | *Thalasseus sandvicensis spp. acuflavidus* | Brazil | 915 | 828 | 87 | Least concern |
| **Cayenne tern** | *Thalasseus sandvicensis spp. eurygnathus* | Argentina | 1 | 1 | 0 | Least concern |
| **Chilean flamingo** | *Phoenicopterus chilensis* | Argentina | 10 | 10 | 0 | Near threatened |
| **Chilean skua** | *Stercorarius chilensis* | Chile | 17 | 17 | 0 | Least concern |
| **Chimango caracara** | *Daptrius chimango* | Chile | 9 | 9 | 0 | Least concern |
| **Cinnamon teal** | *Spatula cyanoptera* | Chile | 5 | 0 | 5 | Least concern |
| **Cocoi heron** | *Ardea cocoi* | Brazil | 1 | 1 | 0 | Least concern |
| **Common quail** | *Coturnix coturnix* | Chile | 5 | 5 | 0 | Least concern |
| **Common tern** | *Sterna hirundo* | Brazil | 27 | 19 | 8 | Least concern |
| **Coscoroba swan** | *Coscoroba coscoroba* | Brazil | 9 | 9 | 0 | Least concern |
|  |  | Chile | 15 | 12 | 3 |  |
| **Crested caracara** | *Caracara plancus* | Brazil | 2 | 1 | 1 | Least concern |
|  |  | Falkland Is. | 1 | 0 | 1 |  |
| **Dolphin gull** | *Leucophaeus scoresbii* | Chile | 2 | 2 | 0 | Least concern |
| **Elegant tern** | *Thalasseus elegans* | Chile | 11 | 11 | 0 | Near threatened |
| **Emu** | *Dromaius novaehollandiae* | Chile | 1 | 1 | 0 | Least concern |
| **Flightless steamer duck** | *Tachyeres pteneres* | Chile | 4 | 4 | 0 | Least concern |
| **Franklin's gull** | *Leucophaeus pipixcan* | Chile | 5 | 5 | 0 | Least concern |
| **Fregatidae (unidentified)** |  | Ecuador | 3 | 1 | 0 |  |
|  |  | Peru | 12 | 10 | 0 |  |
| **Gentoo penguin** | *Pygoscelis papua* | Falkland Is. | 0 | 2300 | 0 | Least concern |
| **Gray gull** | *Leucophaeus modestus* | Chile | 43 | 43 | 0 | Least concern |
|  |  | Peru | 837 | 831 | 1 |  |
| **Graylag goose** | *Anser anser* | Brazil | 4 | 0 | 0 | Least concern |
|  |  | Chile | 2 | 2 | 0 |  |
|  |  | Uruguay | 2 | 2 | 28 |  |
| **Great black-hawk** | *Buteogallus urubitinga* | Brazil | 1 | 0 | 1 | Least concern |
| **Great egret** | *Ardea alba* | Brazil | 1 | 0 | 1 | Least concern |
|  |  | Peru | 1 | 1 | 0 |  |
|  |  | Uruguay | 1 | 1 | 0 |  |
| **Great frigatebird** | *Fregata minor* | Colombia | 1 | 1 | 0 | Least concern |
|  |  | Ecuador | 1002 | 101 | 0 |  |
| **Great grebe** | *Podiceps major* | Chile | 4 | 4 | 0 | Least concern |
| **Grey-hooded gull** | *Chroicocephalus cirrocephalus* | Brazil | 2 | 0 | 2 | Least concern |
|  |  | Peru | 2 | 2 | 0 |  |
| **Guanay cormorant** | *Leucocarbo bougainvilliorum* | Chile | 161 | 161 | 0 | Near threatened |
|  |  | Peru | 197 | 180 | 1 |  |
| **Gull-billed tern** | *Gelochelidon nilotica* | Argentina | 2 | 2 | 0 | Least concern |
| **Harris's hawk** | *Parabuteo unicinctus* | Chile | 2 | 2 | 0 | Least concern |
| **Helmeted guineafowl** | *Numida meleagris* | Peru | 4 | 1 | 0 | Least concern |
| **House sparrow** | *Passer domesticus* | Chile | 1 | 1 | 0 | Least concern |
| **Humboldt penguin** | *Spheniscus humboldti* | Chile | 5 | 5 | 0 | Vulnerable |
|  |  | Peru | 0 | 1 | 0 |  |
| **Imperial cormorant** | *Leucocarbo atriceps* | Argentina | 2 | 2 | 0 | Least concern |
|  |  | Chile | 3 | 3 | 0 |  |
| **Inca tern** | *Larosterna inca* | Chile | 24 | 24 | 0 | Near threatened |
| **Indian peafowl** | *Pavo cristatus* | Brazil | 4 | 1 | 0 | Least concern |
|  |  | Uruguay | 1 | 1 | 0 |  |
| **Kelp gull** | *Larus dominicanus* | Argentina | 22 | 22 | 0 | Least concern |
|  |  | Chile | 71 | 68 | 3 |  |
|  |  | Peru | 16 | 15 | 1 |  |
| **Laughing gull** | *Leucophaeus atricilla* | Ecuador | 1 | 0 | 0 | Least concern |
| **Lesser horned owl** | *Bubo magellanicus* | Chile | 1 | 1 | 0 | Least concern |
| **Lesser yellowlegs** | *Tringa flavipes* | Chile | 1 | 1 | 0 | Vulnerable |
| **Magnificent frigatebird** | *Fregata magnificens* | Brazil | 1 | 0 | 1 | Least concern |
|  |  | Ecuador | 6000 | 6000 | 0 |  |
|  |  | Peru | 210 | 104 | 0 |  |
| **Mandarin duck** | *Aix galericulata* | Chile | 4 | 0 | 4 | Least concern |
| **Manx shearwater** | *Puffinus puffinus* | Brazil | 4 | 1 | 3 | Least concern |
| **Muscovy duck** | *Cairina moschata* | Brazil | 8 | 8 | 0 | Least concern |
| **Mute swan** | *Cygnus olor* | Brazil | 2 | 2 | 0 | Least concern |
| **Neotropic cormorant** | *Nannopterum brasilianum* | Brazil | 2 | 0 | 2 | Least concern |
|  |  | Chile | 17 | 17 | 0 |  |
|  |  | Peru | 0 | 1 | 0 |  |
| **Peregrine falcon** | *Falco peregrinus* | Chile | 3 | 3 | 0 | Least concern |
|  |  | Peru | 2 | 2 | 0 |  |
| **Peruvian booby** | *Sula variegata* | Chile | 199 | 199 | 0 | Least concern |
|  |  | Peru | 457 | 459 | 6 |  |
| **Peruvian diving petrel** | *Pelecanoides garnotii* | Chile | 2 | 2 | 0 | Near threatened |
| **Peruvian pelican** | *Pelecanus thagus* | Chile | 189 | 154 | 35 | Near threatened |
|  |  | Peru | 707 | 583 | 31 |  |
| **Rallidae (unidentified)** |  | Argentina | 4 | 4 | 0 |  |
| **Red-footed booby** | *Sula sula* | Ecuador | 6 | 5 | 0 | Least concern |
| **Red-gartered coot** | *Fulica armillata* | Chile | 3 | 3 | 0 | Least concern |
| **Red-legged cormorant** | *Poikilocarbo gaimardi* | Chile | 18 | 18 | 0 | Near threatened |
|  |  | Peru | 22 | 21 | 1 |  |
| **Ringed teal** | *Callonetta leucophrys* | Brazil | 2 | 2 | 0 | Least concern |
| **Roadside hawk** | *Rupornis magnirostris* | Brazil | 1 | 0 | 1 | Least concern |
| **Rock pigeon** | *Columba livia* | Peru | 2 | 0 | 2 | Least concern |
| **Royal tern** | *Thalasseus maximus* | Argentina | 3 | 3 | 0 | Least concern |
|  |  | Brazil | 88 | 36 | 52 |  |
|  |  | Uruguay | 1 | 1 | 0 |  |
| **Rufous hornero** | *Furnarius rufus* | Brazil | 1 | 0 | 0 | Least concern |
| **Sanderling** | *Calidris alba* | Chile | 5 | 5 | 0 | Least concern |
| **Slender-billed parakeet** | *Enicognathus leptorhynchus* | Chile | 30 | 24 | 6 | Least concern |
| Snowy egret | *Egretta thula* | Brazil | 1 | 0 | 1 | Least concern |
|  |  | Chile | 3 | 3 | 0 |  |
| Sooty shearwater | *Ardenna grisea* | Chile | 3 | 3 | 0 | Near threatened |
|  |  | Peru | 6 | 1 | 0 |  |
| South American tern | *Sterna hirundinacea* | Argentina | 208 | 208 | 0 | Least concern |
|  |  | Brazil | 15 | 6 | 9 |  |
|  |  | Chile | 85 | 85 | 0 |  |
|  |  | Uruguay | 5 | 4 | 1 |  |
| South polar skua | *Stercorarius maccormicki* | Peru | 25 | 20 | 5 | Least concern |
| Southern fulmar | *Fulmarus glacialis* | Falkland Is. | 2 | 2 | 0 | Least concern |
| Southern giant-petrel | *Macronectes giganteus* | Chile | 2 | 2 | 0 | Least concern |
| Southern lapwing | *Vanellus chilensis* | Brazil | 1 | 0 | 1 | Least concern |
|  |  | Chile | 2 | 2 | 0 |  |
| Southern rockhopper penguin | *Eudyptes chrysocome* | Falkland Is. | 0 | 30 | 0 | Vulnerable |
| Spheniscidae (unidentified) |  | Falkland Is. | 0 | 400 | 0 |  |
| Strigiformes (unidentified) |  | Peru | 1 | 1 | 0 |  |
| Tawny eagle | *Aquila rapax* | Peru | 3 | 1 | 0 | Vulnerable |
| Tropical screech-owl | *Megascops choliba* | Brazil | 1 | 0 | 1 | Least concern |
| Turkey vulture | *Cathartes aura* | Chile | 31 | 30 | 1 | Least concern |
| Upland goose | *Chloephaga picta* | Chile | 7 | 6 | 1 | Least concern |
| Variable hawk | *Geranoaetus polyosoma* | Falkland Is. | 1 | 1 | 0 | Least concern |
| Waved albatross | *Phoebastria irrorata* | Peru | 4 | 4 | 0 | Critically endangered |
| Whimbrel | *Numenius phaeopus* | Chile | 2 | 2 | 0 | Least concern |
|  |  | Peru | 1 | 0 | 1 |  |
| White-chinned petrel | *Procellaria aequinoctialis* | Brazil | 2 | 0 | 2 | Vulnerable |
| **White-faced ibis** | *Plegadis chihi* | Brazil | 11 | 10 | 0 | Least concern |
| **White-faced whistling duck** | *Dendrocygna viduata* | Brazil | 16 | 0 | 0 | Least concern |
| **Wilson's storm-petrel** | *Oceanites oceanicus* | Chile | 1 | 1 | 0 | Least concern |
| **Wood Duck** | *Aix sponsa* | Brazil | 1 | 1 | 0 | Least concern |
|  |  | Chile | 24 | 0 | 24 |  |
| **Yellow-billed pintail** | *Anas georgica* | Chile | 7 | 7 | 0 | Least concern |
| **Yellow-billed teal** | *Anas flavirostris* | Chile | 11 | 11 | 0 | Least concern |
| **Yellow-crowned night heron** | *Nyctanassa violacea* | Ecuador | 1 | 1 | 0 | Least concern |
| **TOTAL** |  |  | **13118** | **14144** | **361** |  |

In terms of total cases and deaths, Ecuador reported the highest numbers, with 7,017 confirmed cases and 6,111 fatalities, followed by Peru, Brazil, and Chile (Figure 2). Most of Ecuador’s cases were associated with the magnificent frigatebird (*Fregata magnificens*), which accounted for over 6,000 cases and deaths. In Peru, the species most affected was the gray gull (*Leucophaeus modestus*), in Chile the Peruvian booby (*Sula variegata*), and in Brazil the Cabot’s tern (*Thalasseus sandvicensis* spp. *acuflavidus*).

Among the wild bird species reported with HPAI in South America, 27 (24.77%) belonged to the order Charadriiformes, followed by Anseriformes (n=17, 15.6%), Suliformes (n=12, 11%), and Procellariiformes (n=10, 9.17%) (Figure 3). However, when considering the impact in terms of the number of cases and mortality, Suliformes were the most affected order, mainly due to the magnificent frigatebird. The second most affected order was *Sphenisciformes*, largely driven by the high mortality observed in Gentoo penguins (*Pygoscelis papua*) in the Falkland Islands. Charadriiformes and Pelecaniformes ranked third and fourth, respectively, in terms of overall impact (Figure 4). Notably, during the recurrence event reported by Brazil beginning in May 2025, 10 of the 12 affected species were Anseriformes.

**Figure 3.**
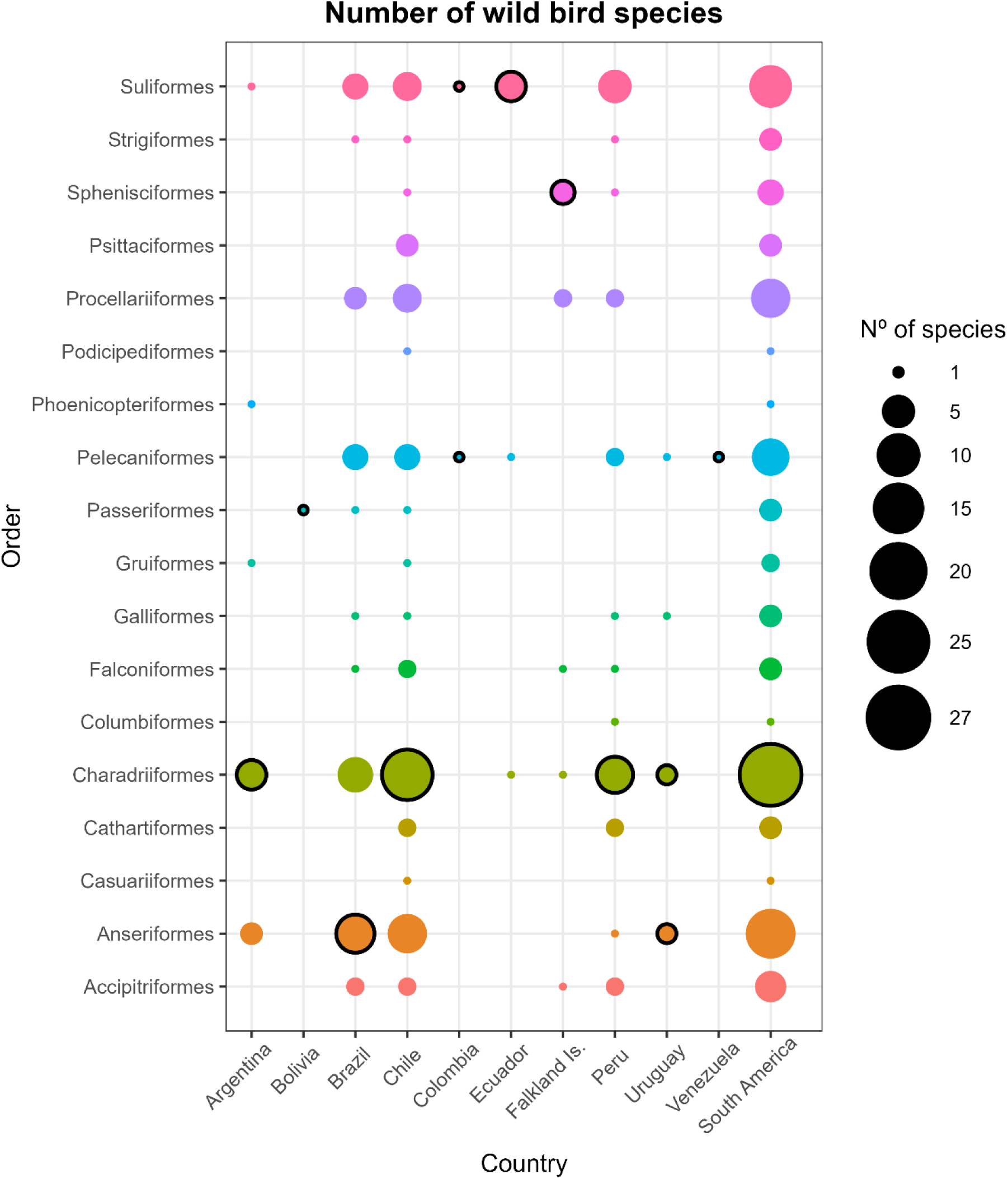
Number of species affected by H5N1 per order per South American country reported to the World Organization for Animal Health (WOAH) until the 30^th^ of May 2025, retrieved from the World Animal Health Information System (WAHIS).

**Figure 4.**
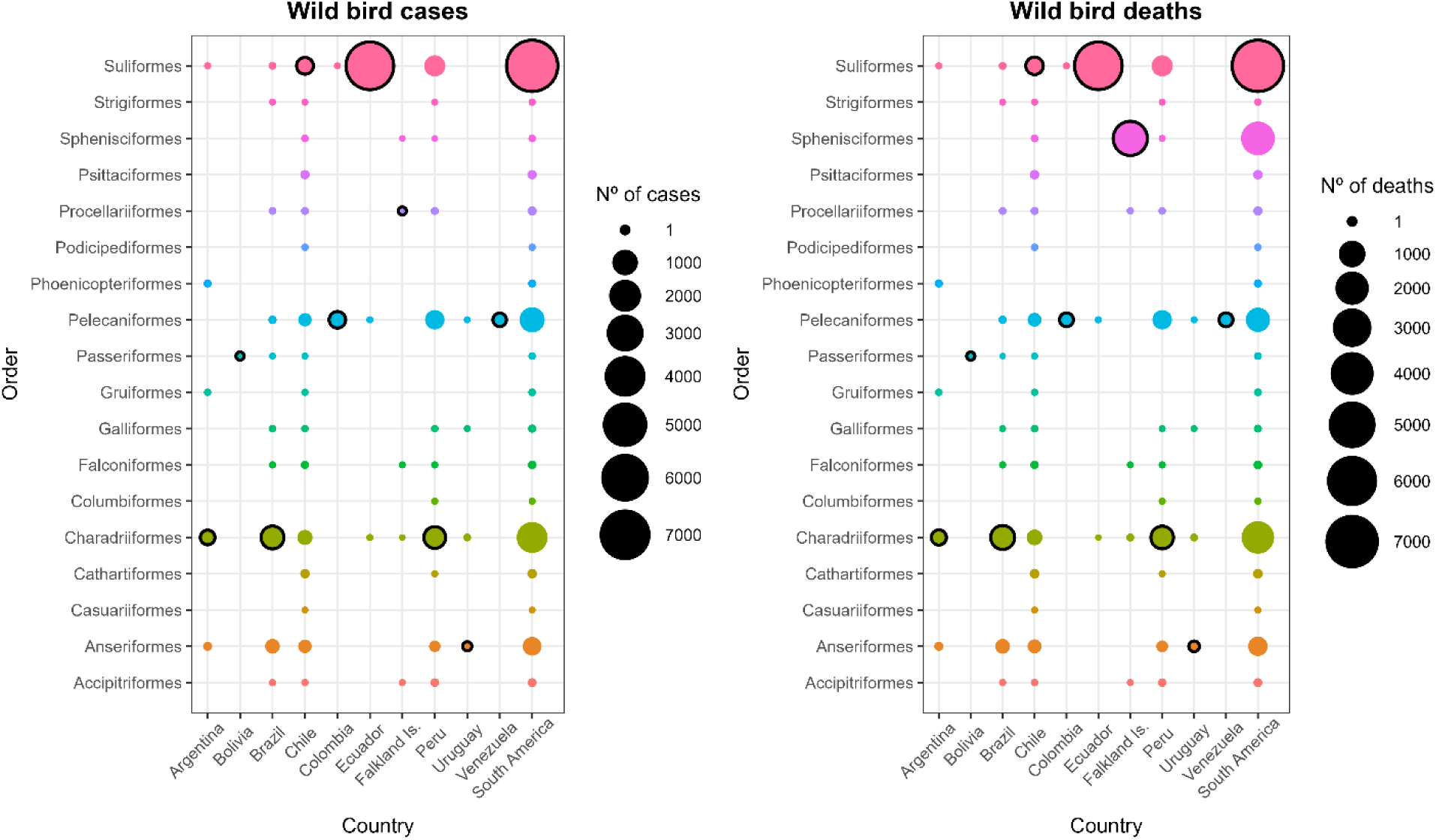
Number of H5N1 cases and deaths (including eliminations) per order per South American country reported to the World Organization for Animal Health (WOAH) until the 30^th^ of May 2025, retrieved from the World Animal Health Information System (WAHIS).

The most affected wild bird species varied by country. In Argentina, the South American tern (*Sterna hirundinacea*) accounted for the highest number of cases, while in Bolivia, the only species reported was the blue-and-white swallow (*Pygochelidon cyanoleuca*). In Brazil, the most affected species was Cabot’s tern (*Thalasseus sandvicensis* spp. *acuflavidus*), and in Chile, the Peruvian booby (*Sula variegata*). Colombia reported the brown pelican (*Pelecanus occidentalis*) as the most affected species, as did Venezuela, where it was the only species identified. In Ecuador, the most impacted species was the magnificent frigatebird (*Fregata magnificens*), while in the Falkland Islands, it was the Gentoo penguin (*Pygoscelis papua*). The grey gull (*Leucophaeus modestus*) was predominant in Peru, and the black-necked swan (*Cygnus melancoryphus*) in Uruguay (Table 1).

Notably, a considerable proportion of affected species were shared across multiple countries. Of the 104 identified wild bird species reported with HPAI, 33.65% (n=35) were detected in more than one country. Among these, the South American tern (*Sterna hirundinacea*) and the black-necked swan (*Cygnus melancoryphus*) were each reported in four countries. Peru exhibited the highest proportion of shared species, with 81.48% (22/27) of its affected species also reported in other countries, 16 of them shared with Chile. In turn, Chile reported 42.11% (24/57) of its species in common with other countries, primarily with Peru, while Brazil shared 41.17% (14/34), mostly with Chile.

Among the identified affected species, 59.62% (62/104) of the wild bird species reported as HPAI H5N1 positive in South America are considered migrants. Within this group, 20 species (32.26%) were classified as trans-equatorial migrants, and 42 (67.74%) as non-trans-equatorial migrants (Supplementary Table 2).

Most of the wild birds affected by HPAI in South America are classified as Least Concern by the IUCN (Table 1). However, eight species are classified as Near Threatened and six as Vulnerable. Out of the six species classified as Vulnerable, two were in captivity. The tawny eagle (*Aquila rapax*) was reported from the Zoológico de la Municipalidad Distrital de Sucre in Peru, and the Andean condor (*Vultur gryphus*) was reported in the Zoológico Las Rocas in Peru. Only one species was classified as Critically Endangered, the Waved albatross (*Phoebastria irrorata*), which was found in the wild in Chile, where only four individuals were affected.

#### Wild mammals

Up to May 2025, no cases of H5N1 have been reported in domestic mammals in South America, however, infections have been documented in wild mammals. A total of 1,141 confirmed cases and 5,772 deaths/eliminations have been reported. As shown in Table 2, the most affected species are marine mammals. Exceptions include Geoffroy’s cat (*Leopardus geoffroyi*), detected in the wild in Chile, a lion (*Panthera leo*) at the Huancayo Zoo in Peru, and a South American coati (*Nasua nasua*) at the Tálice Ecopark in Uruguay. It is important to note that lions are not native to Peru or any South American country, whereas the South American coati is native to Uruguay and other regions of the continent.

**Table 2.** Wild mammal species that were reported by South American countries to the World Organization for Animal Health (WOAH) for “Influenza A viruses of high pathogenicity” until the 30^th^ of May 2025, which are present in the World Animal Health Information System (WAHIS). The global assessment of the IUCN Red List Category was included for each species.

| Species | Scientific name | Country | Nº of cases | Nº of deaths | Nº of eliminated | IUCN Assessment |
| --- | --- | --- | --- | --- | --- | --- |
| Geoffroy's Cat | *Leopardus geoffroyi* | Chile | 1 | 1 |  | Least concern |
| Lion | *Panthera leo* | Peru | 1 | 1 |  | Vulnerable |
| Marine otter | *Lontra felina* | Chile | 2 | 2 |  | Endangered |
| Southern river otter | *Lontra provocax* | Chile | 1 | 1 |  | Endangered |
| South American Coati | *Nasua nasua* | Uruguay | 16 | 16 |  | Least concern |
| South American fur seal | *Arctocephalus australis* | Argentina |  | 44 |  | Least concern |
|  |  | Uruguay | 8 | 8 |  |  |
|  |  | Brazil | 439 | 381 | 1 |  |
| South American sea lion | *Otaria flavescens* | Peru | 2 | 2 |  | Least concern |
|  |  | Chile | 36 | 36 |  |  |
|  |  | Argentina | 17 | 2134 |  |  |
|  |  | Uruguay | 32 | 32 |  |  |
|  |  | Brazil | 578 | 488 | 6 |  |
| Southern elephant seal | *Mirounga leonina* | Argentina | 8 | 2579 |  | Least concern |
|  |  | Falkland Islands |  | 40 |  |  |
| **TOTAL** |  |  | **1141** | **5765** | **7** |  |

The most reported affected marine mammal species was the South American sea lion (*Otaria flavescens*), with reported cases in five South American countries. It was followed by the Southern elephant seal (*Mirounga leonina*), which was reported only in Argentina and the Falkland Islands. Argentina documented the highest number of reported wild mammals, followed by Brazil (Supplementary Table 1). Among the species reported, only the southern elephant seal is considered migratory according to the IUCN Red List (Table 2). The most endangered mammal species affected in South America were the southern river otter (*Lontra provocax*) and the marine otter (*Lontra felina*), both reported in Chile and currently classified as Endangered. In contrast, the remaining wild mammal species affected by H5N1 are listed as Least Concern and represented most reported cases. All wild mammal species infected with the virus belong to the order *Carnivora*.

#### Presence of HPAI in Antarctica

The first report of HPAI was on the 7^th^ of October 2023, in the South Georgia and the South Sandwich Islands (SGSSI), where four brown Skua (*Stercorarius antarcticus*) tested positive for H5N1 in Bird Island. Later, H5N1 was found in the South Polar Skua (*Stercorarius maccormicki*) in the Argentinean Antarctic base “Primavera” in January 2024.

The wild bird species reported in the Antarctica region can be found on Table 3, all the species are of the order Charadriiformes. The most affected bird was the brown skua, with 14 cases and deaths in the SGSSI. The second most reported bird was the kelp gull (*Larus dominicanus*), which was also reported in Argentina, Chile, and Peru. The brown skua and the south polar skua are considered migrants, while the kelp gull is not migratory.

**Table 3.** Wild bird species in the Antarctica region that were reported by South American countries to the World Organization for Animal Health (WOAH) for “Influenza A viruses of high pathogenicity” until the 30^th^ of May 2025, which are present in the World Animal Health Information System (WAHIS). The global assessment of the IUCN Red List Category was included for each species.

| Species | Scientific name | Antarctic region | Nº of cases | Nº of deaths | Nº of eliminated | IUCN Assessment |
| --- | --- | --- | --- | --- | --- | --- |
| Brown skua | *Stercorarius antarcticus* | Lagoon Island | 2 | 2 | 0 | Least concern |
|  |  | South Georgia and the South Sandwich Islands | 14 | 14 | 0 |  |
| South polar skua | *Stercorarius maccormicki* | Antarctic base “Primavera” | 1 | 1 | 0 | Least concern |
| Kelp gull | *Larus dominicanus* | South Georgia and the South Sandwich Islands | 8 | 8 | 0 | Least concern |
| TOTAL |  |  | 25 | 25 | 0 |  |

#### Phylogenetic analysis of available sequences

The maximum likelihood phylogeny of the HA segment of HPAI H5N1, constructed with a dataset of 130 viral sequences from South America together with representative isolates from North America, revealed a strong clustering by genotype and geography (Figure 5). All Chilean sequences grouped within the B3.2 genotype, forming a monophyletic clade that also includes viruses from Brazil, Peru, Uruguay, Argentina, and Bolivia, and that is clearly distinct from other South American lineages such as B1.2 and B2.2 detected in Colombia and Venezuela. Within Chile, viruses sampled from wild birds, marine mammals, and domestic poultry showed very limited genetic divergence, consistent with a recent introduction event followed by local dissemination across host species. A similar pattern was observed in most South American countries, where a single genotype was detected, predominantly B3.2. Conversely, the presence of mixed clades containing sequences from different countries and host species suggests multiple independent introduction events and potential transboundary spread. Notably, several clades included sequences from both domestic and wild animals, providing evidence for interspecies transmission events (spillover) North America, however, displayed greater genetic heterogeneity, with the co-circulation of multiple genotypes, highlighting a more complex evolutionary and epidemiological scenario. Altogether, this phylogenetic structure indicates that H5N1 circulation in Chile, and more broadly in South America, is dominated by a single genotype (B3.2), with repeated evidence of interspecies transmission but without signs of multiple introductions within each country.

**Figure 5.**
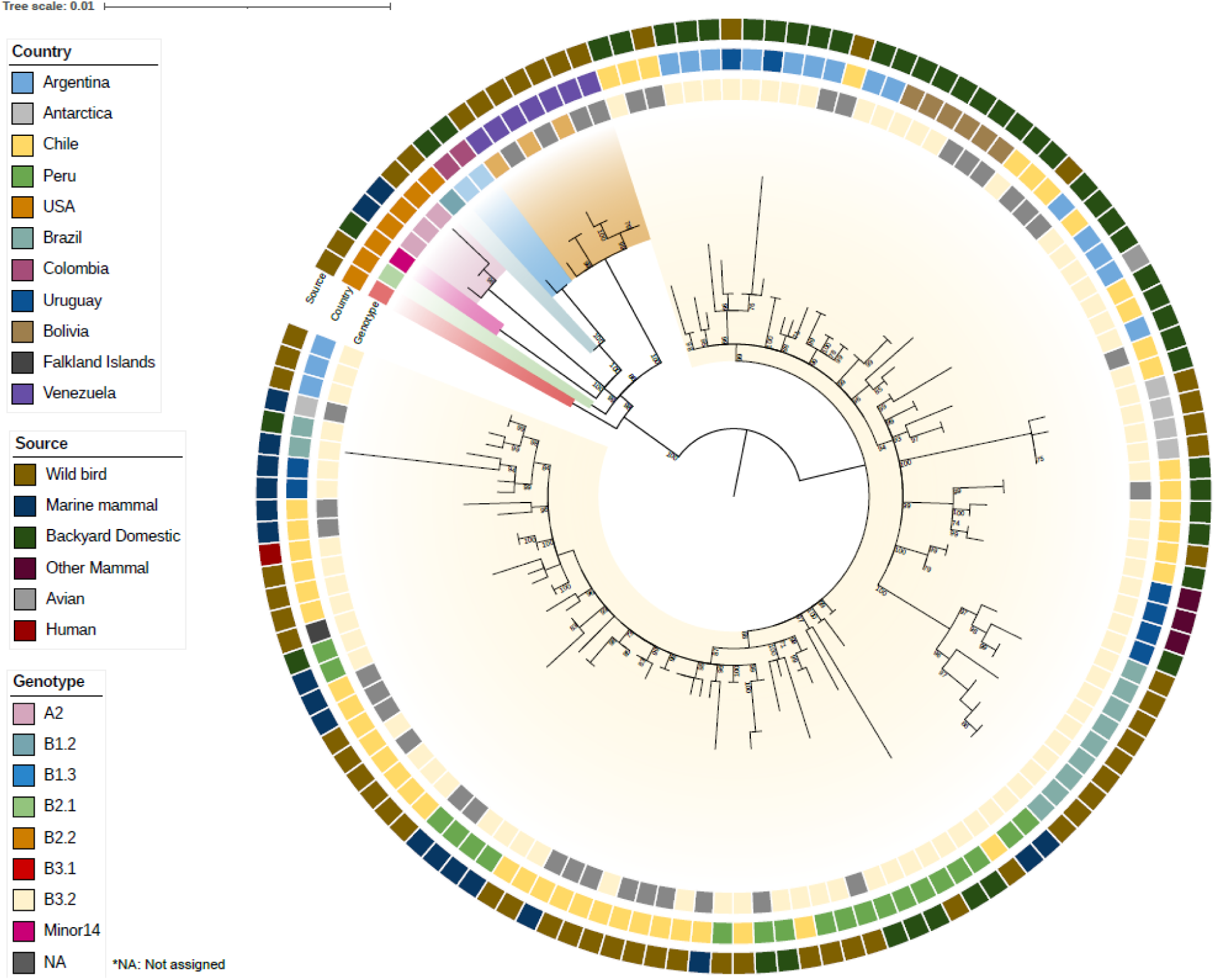
Maximum likelihood phylogenetic tree based on HA segment sequences of H5N1 2.3.4.4b viruses. The tree includes viruses from various host species (wild birds, marine mammals, domestic animals, other mammals and human) collected across multiple South American countries. The concentric rings from the center outward indicate viral genotype, ecological source of the virus, and country of origin.

The time-scaled phylogeny (Figure S1) further resolved the timing and number of introduction events of HPAI H5N1 into South America during 2022. In this analysis, the stem age of a clade (represented by a circle) was defined as the divergence between the focal lineage and its closest outgroup, representing the earliest possible timing of introduction into a region or host. The crown age (represented by a triangle) corresponded to the time to the most recent common ancestor (tMRCA) of all sequences within the clade, marking the latest point by which the lineage was already established and diversifying. Thus, the true introduction is expected to have occurred along the branch between the stem and crown ages, which we report as the estimated introduction window.

Three independent introductions of HPAI H5N1 into South America were identified. The first was traced to Colombia (red node), with a stem age of 2022.1146 (95% HPD: 2021.8771– 2022.1944) and a crown age of 2022.765 (95% HPD: 2022.6501–2022.8312), placing the arrival between the first and third quarters of 2022. A second introduction occurred in Venezuela (green node), with a stem age of 2022.1057 (95% HPD: 2021.8777–2022.2069) and a crown age of 2022.7544 (95% HPD: 2022.8498–2022.8631), indicating a nearly synchronous entry. A third introduction seeded two divergent lineages within South America, with the split dated to mid- 2022 (yellow node). Both sublineages reached Chile almost simultaneously: the first (blue node) with a stem age of 2022.1607 (95% HPD: 2021.9691–2022.2929) and a crown age of 2022.8927 (95% HPD: 2022.8498–2022.9290), and the second (light-blue node) with a crown age of 2022.9061 (95% HPD: 2022.8271–2022.9452), consistent with introductions occurring during the last quarter of 2022. Cross-species transmission to marine mammals was detected shortly thereafter. The earliest spillover event into marine mammals (purple node) was dated to late 2022–early 2023, with a stem age of 2022.89 (95% HPD: 2022.837–2022.9508) and a crown age of 2023.0495 (95% HPD: 2023.0161–2023.0658). A second mammalian clade (light purple node) emerged almost simultaneously, with a crown age of 2023.0332 (95% HPD: 2022.9793–2023.0603), supporting the occurrence of repeated independent spillover events into marine mammals. Finally, the first human case in Chile (orange node) clustered within the Chile–Peru coastal clade, with the node preceding the human sequence dated to early 2023 (stem age 2023.037; 95% HPD: 2022.9241–2023.1372) and the human sequence itself representing the crown age at 2023.1862 (95% HPD: 2023.1298–2023.2247).

## DISCUSSION

The findings of this study provide a comprehensive overview of the impact of the H5N1 avian influenza virus on domestic bird and wildlife populations in South America, highlighting significant ecological and economic implications. We found that more than 8.8 million domestic birds, backyard and commercial, died or were eliminated due to H5N1 in South America.

### HPAI in domestic birds and the need to strengthen backyard poultry surveillance

The countries most affected by H5N1 in backyard poultry were Argentina, Chile, Colombia and Peru, while commercial poultry losses were highest in Argentina, Chile, and Ecuador. According to FAO data, Brazil is the largest producer of domestic birds in South America, with over 6.5 billion birds in 2023, followed by Colombia (1.09 billion), Peru (818 million), and Argentina (803 million) (48). Notably, Brazil reported H5N1 only in backyard birds, with no cases in commercial operations. This suggests that Brazil’s biosecurity and surveillance measures in the commercial poultry sector may be particularly effective and could serve as a model for other countries in the region. In contrast, the next three largest producers, Colombia, Peru, and Argentina, experienced significant losses in both backyard and commercial birds, raising concerns about potential impacts on food availability at national and regional levels. Smaller producers such as Bolivia, Chile, and Ecuador experienced proportionally higher losses. Ecuador, with 287 million domestic birds (48), was the most affected country in terms of commercial poultry, reporting over 1.2 million cases.

Most domestic bird cases occurred in commercial settings, likely reflecting more active surveillance due to the sector’s economic importance. However, backyard poultry remain more vulnerable to infection. In six of the eleven countries that reported outbreaks, the virus was first detected in wild birds and subsequently in backyard flocks. These birds are typically kept for self-consumption, often under poor biosecurity conditions and with frequent contact with wildlife (49), creating an environment conducive to viral reassortment (50). Strengthening surveillance in backyard systems is essential, particularly in countries with large rural populations such as Ecuador, Bolivia, and Paraguay (48).

### Widespread mortality and conservation threats among South America’s wild birds

Wild birds in South America experienced severe mortality due to HPAI H5N1. In Ecuador alone, 6,000 magnificent frigatebirds died. Partners in Flight estimated a population of 130,000 globally; therefore, this means that around 4.8% of the global population of magnificent frigatebirds died due to this virus. This species is capable of migrating from South America to Central America, raising concerns about the potential spread to other colonies and species (32).

Official reports to WOAH likely underestimate the true toll. For example, Chile recorded 94,000 wild bird deaths between December 2022 and 2023 (52), far exceeding the 1,221 cases reported to WOAH. Furthermore, more than 3,700 Humboldt penguins were found stranded in Chile in 2023 and 2024 (53). Similarly, in Peru, Gamarra-Toledo, et al., 2023 (54) documented over 100,000 wild bird deaths across 24 species with signs consistent with H5N1, while official reports listed only 2,374 cases. This discrepancy suggests that monitoring in protected wild areas could enhance surveillance efforts.

Most affected wild bird species are classified as Least Concern, yet species with more vulnerable conservation statuses have also been impacted. The Andean condor, a Vulnerable species native to several South American countries, was reported infected in captivity in Peru. If wild populations become infected, mortality could threaten their long-term survival. A similar situation occurred with the critically endangered California condor (*Gymnogyps californianus*) in the United States, prompting a vaccination campaign to avoid extinction (55). A comparable strategy may become necessary in South America.

The only Critically Endangered species affected was the waved albatross (*Phoebastria irrorata*), with four cases reported in Chile. Found only in Chile, Peru, Colombia, and Ecuador, this species has an extremely limited breeding range, restricted to the Galápagos and La Plata Islands in Ecuador (56). Conservation of the Waved Albatross should prioritize HPAI prevention, as Ecuador—its sole breeding location—has the highest wild bird mortalities in South America. Their migratory behavior also exposes them to infection elsewhere, making coordinated action by Ecuador, Peru, and Chile essential to prevent further declines.

Among HPAI-positive wild birds, 59.62% are migratory species. Trans-equatorial migrants pose the greatest risk for the introduction and spread of avian influenza viruses, with migrations shown to be linked to viral movements (57, 58). Transatlantic migrants were responsible for introducing H5N1 clade 2.3.4.4b from Northwestern Europe to North America via Iceland, Greenland/Arctic, or a direct pelagic route (25). Similarly, the HPAI H7N3 virus detected in Chile was also suggested to have been introduced by wild birds (59). Non-trans-equatorial migrants represented 67.74% of the total migratory birds reported, most performing austral migrations, although some species undertake multiple migration types (32, 33). The southern giant-petrel (*Macronectes giganteus*), for instance, has been recorded migrating north from Antarctica to Brazil (60). Moreover, individuals banded near the Antarctic Peninsula have been found across Australia, South America, South Africa, New Zealand, Madagascar, the Indian Ocean, and the East Sea (61), consistent with juveniles migrating circumpolarly to the northeast while adults stay nearer to breeding sites (61, 62). Although only two individuals tested positive for H5N1 in Chile, this species could spread the virus across multiple continents, including Antarctica.

### HPAI spillover into mammals and implications for surveillance

Up to May 2025, no cases of HPAI H5N1 have been reported in domestic mammals in South America. However, the situation in North America highlights the potential risk, as the United States has confirmed infections with the Eurasian lineage goose/Guangdong clade 2.3.4.4b in dairy cattle, cats, camelids, goats, and swine, while Canada has reported cases in cats and dogs (29). Given the zoonotic potential of these viruses, domestic mammals should be included in South American surveillance strategies. Globally, from January 2003 to April 2025, 973 human cases have been confirmed, with a 48.3% case fatality rate (63), including infections linked to dairy cattle in the United States (64, 65). In South America, only two human cases have been reported: a 9-year-old girl in Ecuador exposed to backyard poultry (66, 67) and a 53-year-old man in Chile, likely infected via environmental exposure near H5N1-positive wild birds (68). In both cases, all close contacts tested negative for influenza.

In contrast, wild mammals in South America have experienced significant mortality. More than 5,772 deaths or eliminations have been reported officially, though actual numbers are likely much higher. In Peru, over 5,000 South American sea lions died between January and April 2023, many with respiratory and neurological symptoms and in areas with H5N1-positive wild birds (69). In Chile, strandings of sea lions rose 1,983% during the first half of 2023, reaching over 22,000 individuals by 2024 (53, 70). Chile also detected H5N1 in Chilean dolphins and Burmeister’s porpoises, with numerous strandings of marine otters, dolphins, and porpoises (53). Argentina reported over 17,000 deaths of southern elephant seals at Peninsula Valdés by late 2023, representing a 96% mortality rate (71).

The affected marine otter and the southern river otter, both with limited geographic ranges in South America, are of particular conservation concern (31). A lion infected in a Peruvian zoo, along with previous detections in captive big cats globally, (29, 72) highlights the risk HPAI H5N1 poses to captive wildlife. In Chile, H5N1 was detected in a Geoffroy’s cat found in an area with abundant wild bird mortality (73), and a similar case was reported in the USA (29, 72). These findings underscore the need for enhanced surveillance and research on susceptibility and risk factors in wild felids.

Several wild mammal species affected by H5N1 in North America are also present in South America, including pumas, feral cats, house mice, and American minks, the latter two being introduced species (29, 31). Alien species may act as reservoirs or amplifiers of the virus. Invasive species may aid in the introduction and amplification of pathogens (74), suggesting the need to assess their role in South America.

The temporal pattern of mammal infections, occurring after wild bird outbreaks, suggests that initial transmission likely stems from contact with infected birds. Whether transmission among mammals occurs remains under investigation. Some researchers propose ingestion of infected birds as the primary route (69), while others suggest that mammal-to-mammal transmission may be facilitated by adaptive mutations, as observed in sea lions, red foxes, and humans (75–77).

### Ongoing viral circulation highlights One Health surveillance needs

The emergence of human H5N1 infections in South America is closely linked to viral circulation in wild birds and marine mammals. In December 2022, the first outbreak in Peruvian pelicans was detected in northern Chile, where HPAI prevalence in wild birds increased rapidly from <1% in August-September to 12% by December (78). Maximum-likelihood phylogenetic reconstruction demonstrated wild bird isolates clustered closely with clade 2.3.4.4b strains from North America and showed no mammalian adaptation markers.

Building on initial wild bird analyses, Pardo-Roa et al. (2025) (77) used an expanded dataset that included wild birds, poultry, marine mammals, and a human case from Chile to infer a time-scaled phylogeny, revealing multiple inter- and intra-regional dispersal events, including spread to Argentina, Uruguay, and Brazil. They identified a distinct marine mammal clade with PB2 D701N and Q591K substitutions absent in avian lineages, indicating sustained pinniped transmission along over 3,800 km of coastline. Our Bayesian estimates place the first marine mammal spillover to late 2022–early 2023 between and the Chilean human spillover in Chile to early March 2023, highlighting ongoing viral circulation and cross-species transmission risk beyond (77).

Rivetti et al., 2024 (79), conducted a comprehensive phylogeographic assessment of Brazilian H5N1 isolates, revealing dual Pacific and Atlantic dispersal corridors and identifying South American clades that encompass marine mammal sequences and harbor the same PB2 D701N and Q591K adaptations. Their findings underscore the extensive inter-country connectivity and cross-species transmission dynamics of clade 2.3.4.4b viruses. In concordance with this, our analysis delineates mixed-host clades spanning the entire South American continent and corroborates a Pacific origin for the pinniped spillover event. Moreover, by distinguishing three temporally discrete introductions into South America, and by precisely estimating their dates, our study augments the continental phylogeographic framework established by Rivetti et al., 2024 (79), providing enhanced temporal resolution and reinforcing the imperative for integrated One Health surveillance across avian, marine mammal, and human populations.

### Evidence of H5N1 transmission between avifauna in South America and Antarctica

The emergence of HPAI H5N1 in Antarctica in October 2023 represents a critical expansion of the virus into one of the most ecologically sensitive regions on Earth. Before this event, only LPAI subtypes such as H6N8, H11N2, and H5N5 had been detected on the continent (80–82). The initial detections in South Georgia and the South Sandwich Islands (SGSSI), followed by cases in the Falkland Islands, suggest possible introduction via migratory seabirds, particularly the brown skua, which connects both regions through overlapping wintering grounds (32, 83).

Additional detections in January 2024, including the South Polar Skua and the kelp gull, further support viral circulation within the region, potentially facilitated by both migratory and resident species (84, 85). All reported Antarctic cases have involved species of the order *Charadriiformes*, the same order most frequently affected in South America, suggesting possible taxonomic susceptibility that merits further study. Moreover, several species reported with H5N1 in South America, such as the southern giant petrel, gentoo penguin, and black-browed albatross, also occur in Antarctica (32), raising concerns about the continued spread of the virus across polar ecosystems. These findings highlight the need for enhanced surveillance of migratory seabirds and polar avifauna to better understand and mitigate the risks posed by HPAI in the Antarctic region

In conclusion, between 2022 and 2025, HPAI H5N1 clade 2.3.4.4b caused extensive outbreaks across 11 South American countries, resulting in significant mortality among domestic poultry, wild birds, and marine mammals, and marking its unprecedented spread to Antarctica in early 2024. Migratory birds played a key role in long-distance dissemination, while high mortality in marine mammals suggests potential mammal-to-mammal transmission. Environmental stressors, both acute and chronic, can modify migratory behaviors, species distributions, and interspecies interactions (including with humans and domestic animals), thereby influencing AIV transmission. (86–88). Moreover, rising temperatures may adversely affect reproductive success, thereby influencing the long-term conservation of migratory species (86). The combined effects of climate change and limited surveillance, particularly in backyard and marine systems, pose ongoing challenges to outbreak detection and control. These findings highlight the urgent need for a coordinated One Health approach—integrating wildlife, domestic animal, and human health—to strengthen early warning systems, mitigate zoonotic risks, and prevent future outbreaks.

## MATERIALS AND METHODS

### Data gathering and visualization

We gathered all the reports from the World Animal Health Information System (WAHIS) from the World Organization for Animal Health (WOAH) from the “Influenza A viruses of high pathogenicity (Inf. with) (non-poultry including wild birds) (2017-)” and “High pathogenicity avian influenza viruses (poultry) (Inf. with)” from South American countries, up to the 30^th^ of May 2025 (29). We considered all domestic non-poultry as backyard domestic birds, and poultry were separated into backyard and commercial domestic birds. Case numbers are repeated in the deaths and eliminated columns; therefore, they are not to be added. Susceptible wild bird species that were eliminated but were not infected by H5N1 were not included in the study, while susceptible domestic birds (backyard and commercial) that were eliminated as a method of control were included. The conservation status for each affected species, as well as the migratory information for wild mammals, was retrieved from the IUCN Red List of Threatened Species (31), and the migratory information for wild birds was retrieved from Birds of the World from the Cornell Laboratory of Ornithology (32), which resulted in three categories based on bird migratory pattern: trans-equatorial, non-trans-equatorial and non-migratory in South America. We defined trans-equatorial migration as movements in which birds crossed the equator, migrating between South America and Central or North America. The non-trans-equatorial category included all migratory birds that remained within South America, undertaking austral, longitudinal, or altitudinal migrations, as described in Jahn et al. (2020) (33), as well as those that perform circumpolar movements around Antarctica. We analyzed the retrieved data descriptively, while the timeline graph was created in R (version 4.2.1) with the ‘ggplot2’ package, and the maps with the ‘tmap’, ‘sf’, and ‘rnaturalearth’ packages (34–37).

### Phylogenetic analysis

A maximum likelihood phylogenetic tree was constructed to analyze the genetic diversity of Influenza A virus across different host species and countries in South America. To avoid geographic overrepresentation in the phylogenetic analysis, a maximum of 15 HA sequences per country were included. These sequences were randomly selected while ensuring representation across all available host species, including wild birds, domestic animals, and marine mammals when applicable. This approach allowed for a balanced dataset that captures host diversity without biasing the analysis toward countries with higher sequencing efforts. Complete sequences of the HA segment were downloaded from the GISAID database, including isolates from marine mammals, birds, and humans. The sequences were aligned using MAFFT v7.526 (38) and subsequently manually edited in BioEdit (39) to remove ambiguous or poorly aligned regions. The phylogenetic tree was generated using the IQ-TREE v2.4.0 software (40), employing the best-fit evolutionary model selected automatically through ModelFinder (41) and applying the ultrafast bootstrap method (42) with 10,000 replicates to assess branch support. Tree visualization and annotation were performed in iTOL v4 (43), incorporating metadata associated with each sequence, such as country and host species of origin. Additionally, viral genotypes were assigned to each virus using the Genoflu v1.06 tool (44), enabling a phylogenetic interpretation contextualized according to circulating viral lineages.

We further conducted a time-calibrated phylogenetic analysis using BEAST v1.10.5.0 (45). The dataset consisted of time-stamped sequences (decimal year) and was evaluated for molecular clock signal using TempEst v1.5 (46), which revealed a positive correlation between sampling date and root-to-tip divergence, supporting the application of a molecular clock model. Analyses were performed under a GTR substitution model and included both strict and uncorrelated lognormal relaxed clock models. Tree priors were tested under several demographic models, including coalescent constant size, exponential growth, logistic growth, expansion growth, and Bayesian skygrid. The skygrid model with 10 grid points provided the best fit and was selected for final analysis. Three independent Markov chain Monte Carlo (MCMC) runs were executed, each consisting of 50 million generations, sampling every 5,000 generations. Log and tree files were combined after removing 10% burn-in from each run using LogCombiner. The resulting posterior tree distributions were summarized using TreeAnnotator to obtain the maximum clade credibility (MCC) tree. Phylogenetic trees were visualized using TreeViewer v2.2.0 (47).

---

**Supplementary Table 1.**
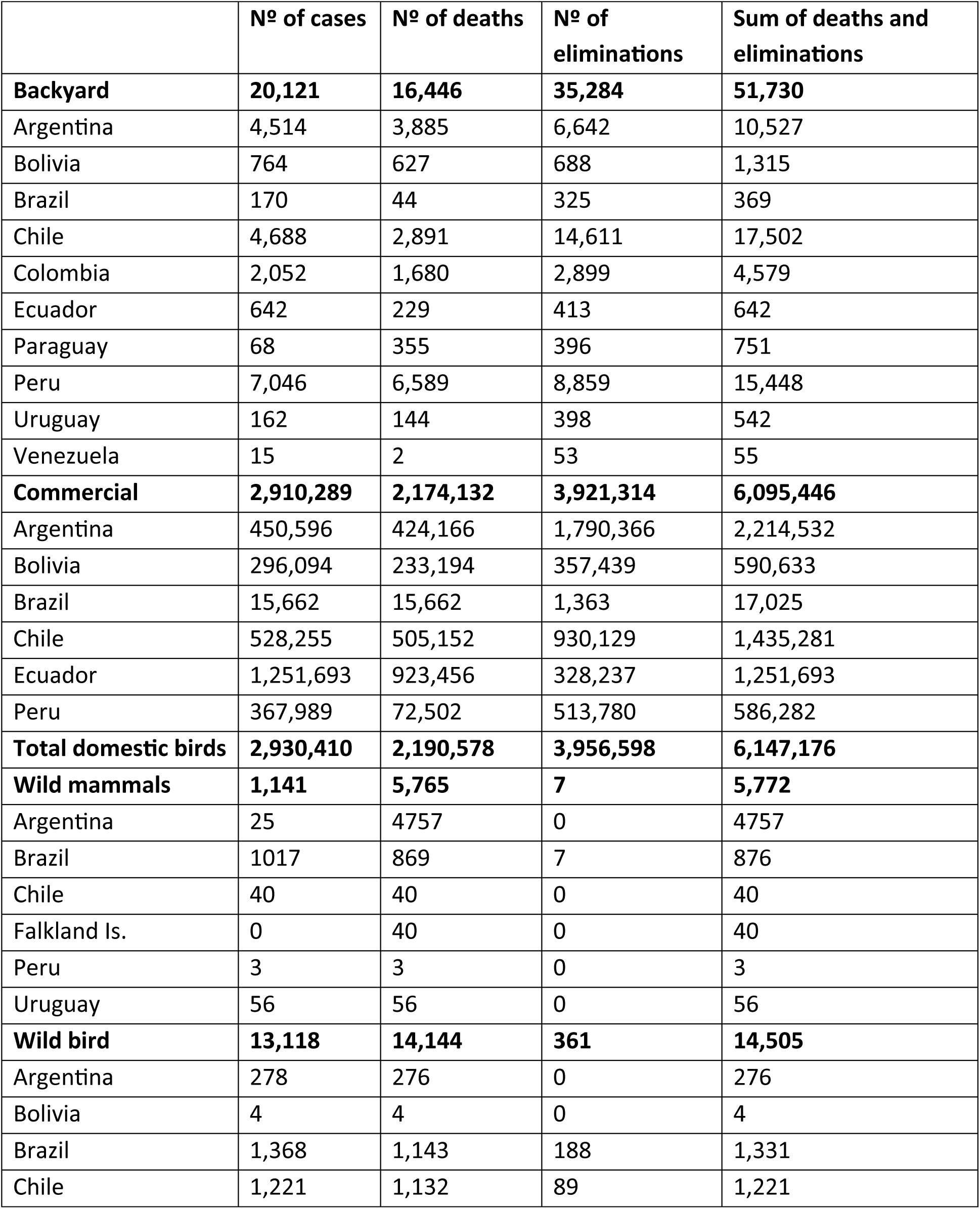
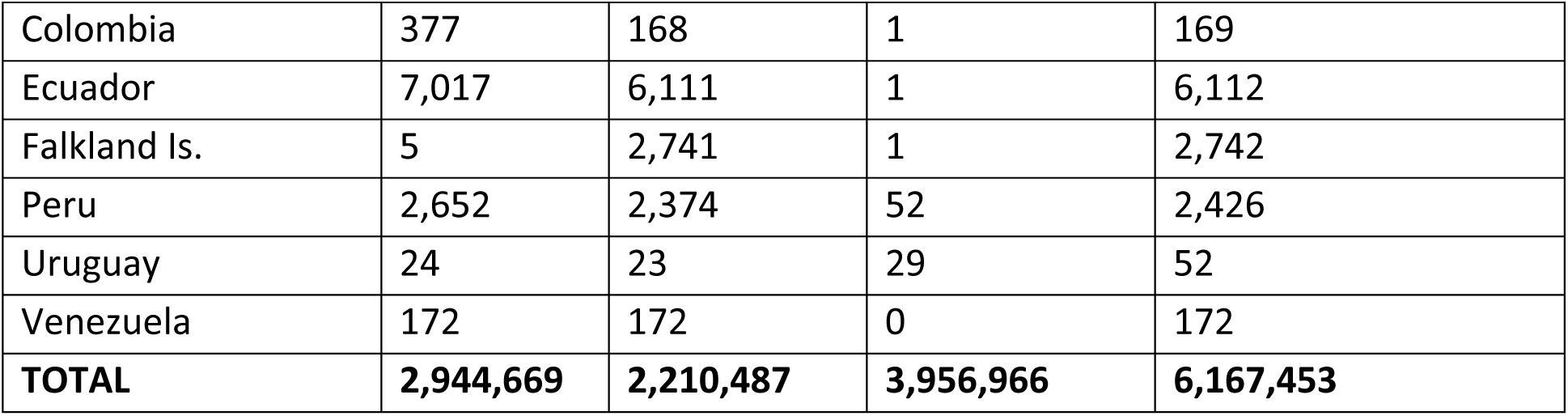
Total positive cases, deaths, and eliminations of host species affected in South American countries, 2022–May 2025.

|  | Nº of cases | Nº of deaths | Nº of eliminations | Sum of deaths and eliminations |
| --- | --- | --- | --- | --- |
| **Backyard** | **20,121** | **16,446** | **35,284** | **51,730** |
| Argentina | 4,514 | 3,885 | 6,642 | 10,527 |
| Bolivia | 764 | 627 | 688 | 1,315 |
| Brazil | 170 | 44 | 325 | 369 |
| Chile | 4,688 | 2,891 | 14,611 | 17,502 |
| Colombia | 2,052 | 1,680 | 2,899 | 4,579 |
| Ecuador | 642 | 229 | 413 | 642 |
| Paraguay | 68 | 355 | 396 | 751 |
| Peru | 7,046 | 6,589 | 8,859 | 15,448 |
| Uruguay | 162 | 144 | 398 | 542 |
| Venezuela | 15 | 2 | 53 | 55 |
| **Commercial** | **2,910,289** | **2,174,132** | **3,921,314** | **6,095,446** |
| Argentina | 450,596 | 424,166 | 1,790,366 | 2,214,532 |
| Bolivia | 296,094 | 233,194 | 357,439 | 590,633 |
| Brazil | 15,662 | 15,662 | 1,363 | 17,025 |
| Chile | 528,255 | 505,152 | 930,129 | 1,435,281 |
| Ecuador | 1,251,693 | 923,456 | 328,237 | 1,251,693 |
| Peru | 367,989 | 72,502 | 513,780 | 586,282 |
| **Total domestic birds** | **2,930,410** | **2,190,578** | **3,956,598** | **6,147,176** |
| **Wild mammals** | **1,141** | **5,765** | **7** | **5,772** |
| Argentina | 25 | 4757 | 0 | 4757 |
| Brazil | 1017 | 869 | 7 | 876 |
| Chile | 40 | 40 | 0 | 40 |
| Falkland Is. | 0 | 40 | 0 | 40 |
| Peru | 3 | 3 | 0 | 3 |
| Uruguay | 56 | 56 | 0 | 56 |
| **Wild bird** | **13,118** | **14,144** | **361** | **14,505** |
| Argentina | 278 | 276 | 0 | 276 |
| Bolivia | 4 | 4 | 0 | 4 |
| Brazil | 1,368 | 1,143 | 188 | 1,331 |
| Chile | 1,221 | 1,132 | 89 | 1,221 |
| Colombia | 377 | 168 | 1 | 169 |
| Ecuador | 7,017 | 6,111 | 1 | 6,112 |
| Falkland Is. | 5 | 2,741 | 1 | 2,742 |
| Peru | 2,652 | 2,374 | 52 | 2,426 |
| Uruguay | 24 | 23 | 29 | 52 |
| Venezuela | 172 | 172 | 0 | 172 |
| **TOTAL** | **2,944,669** | **2,210,487** | **3,956,966** | **6,167,453** |

**Supplementary Table 2.**
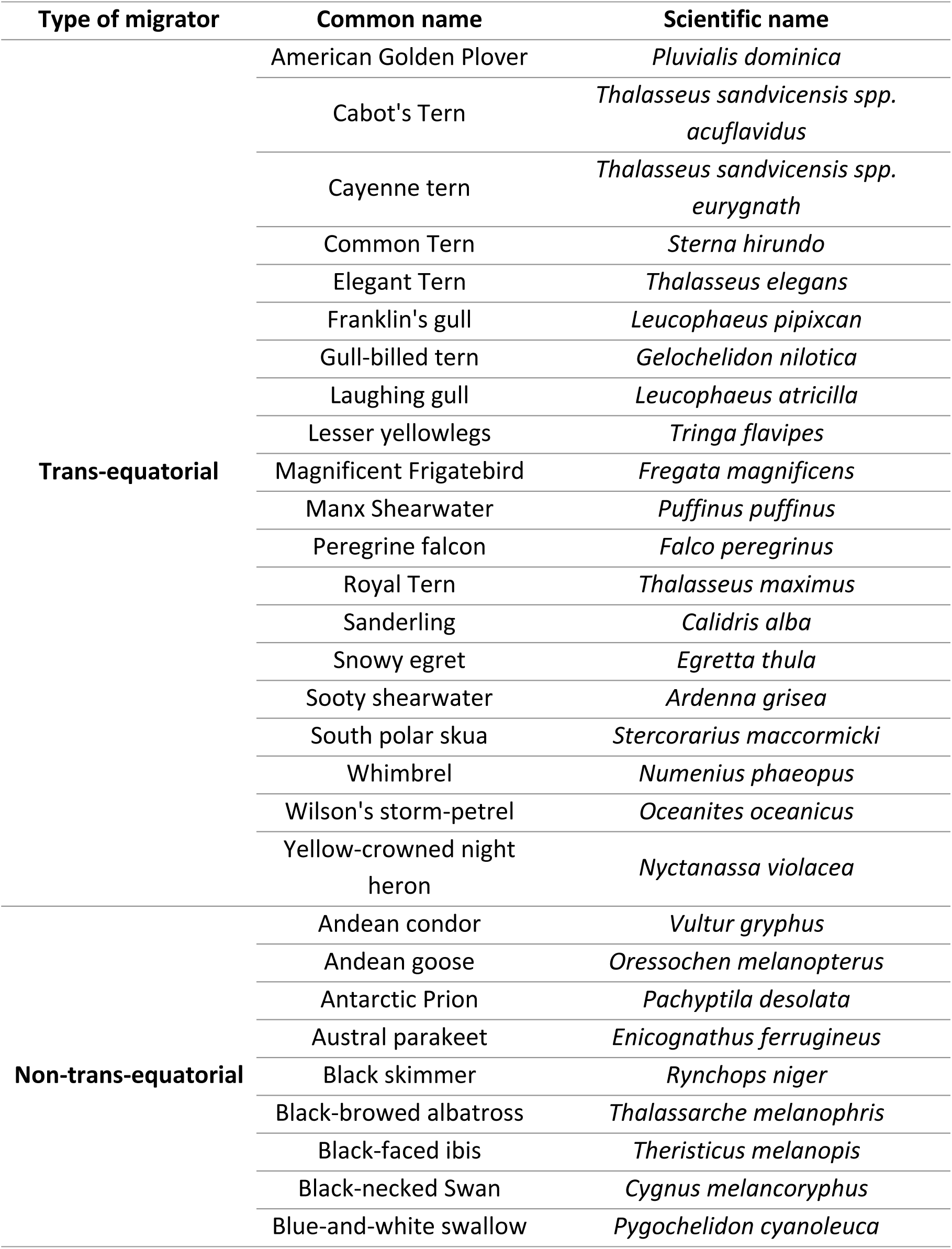
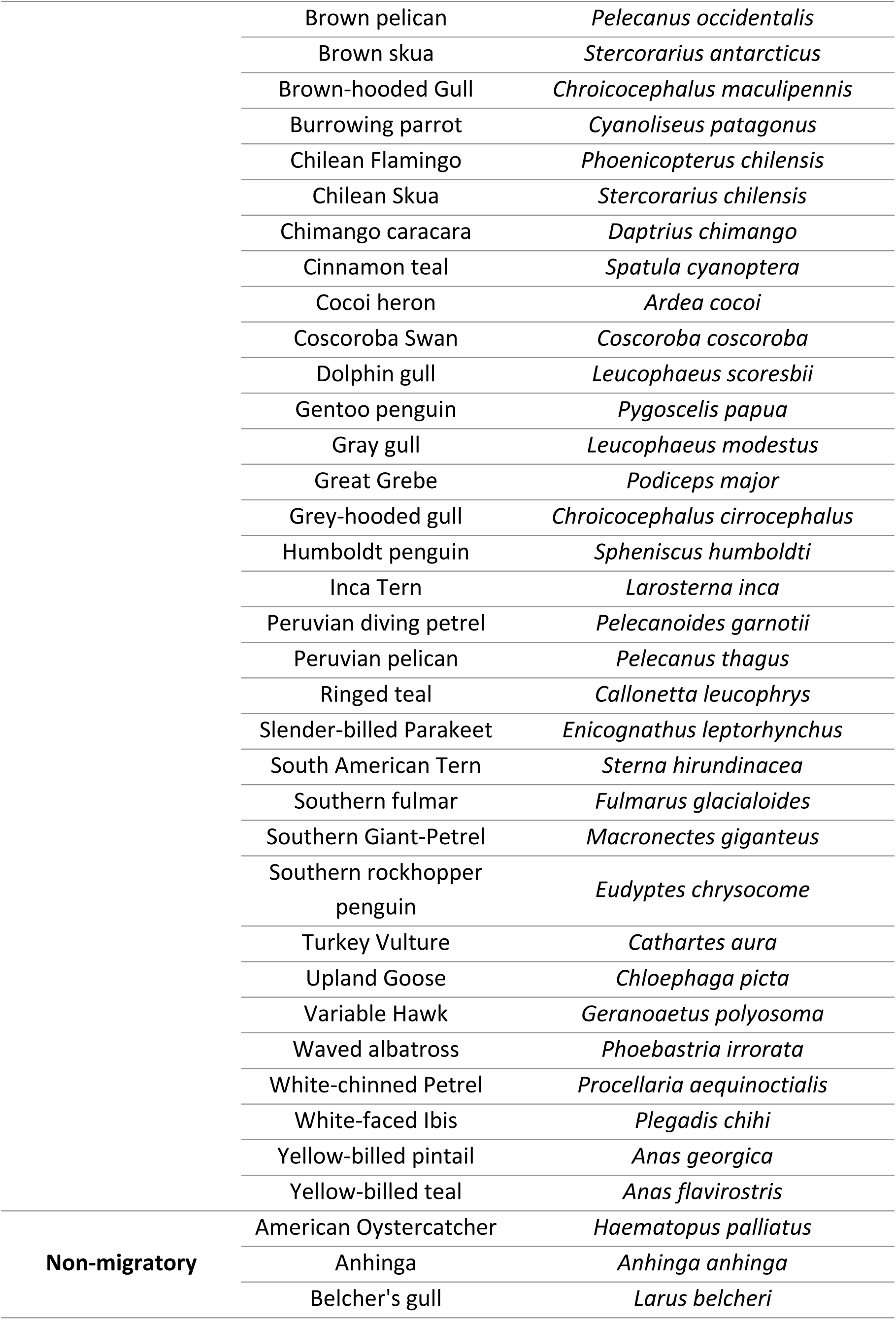
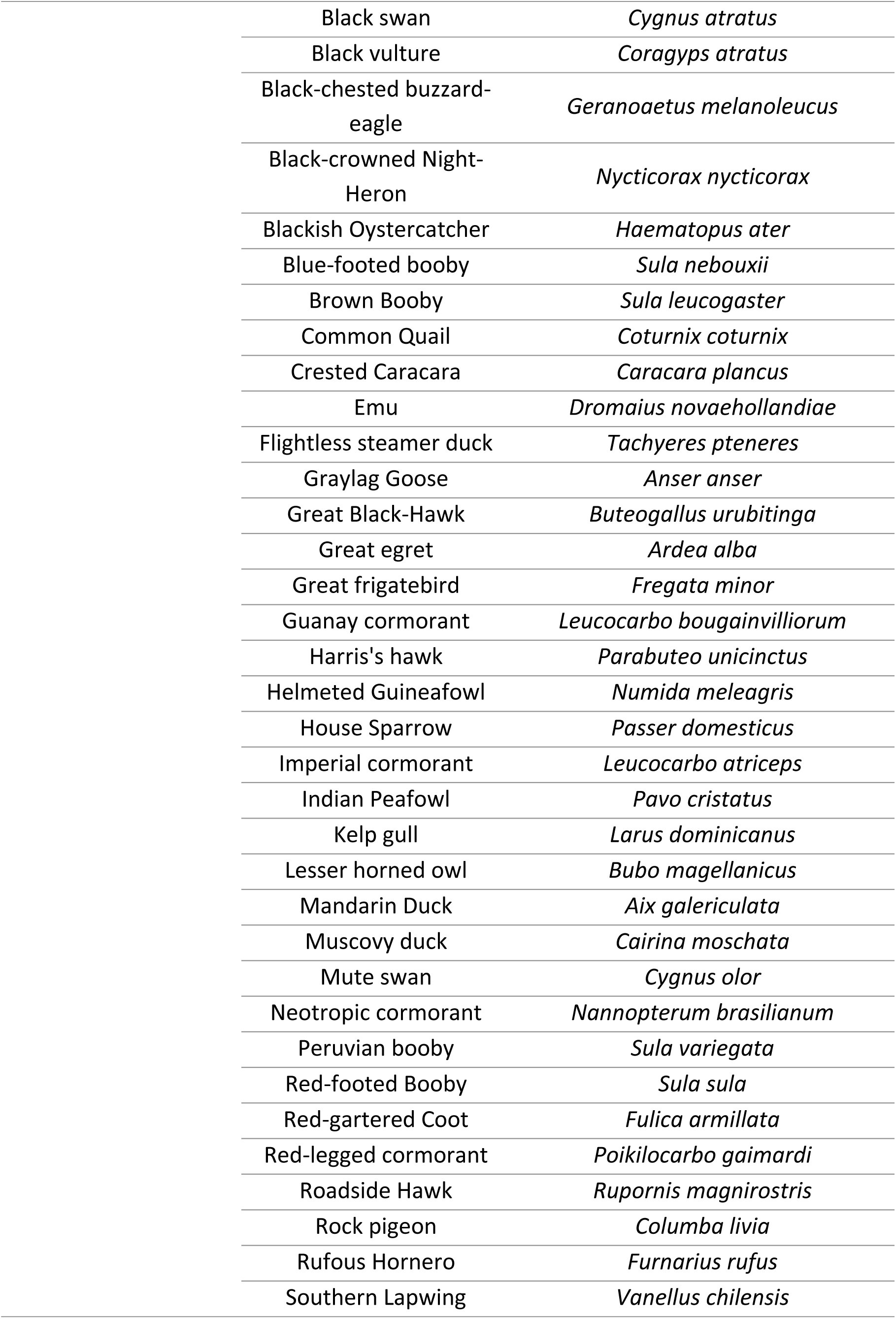
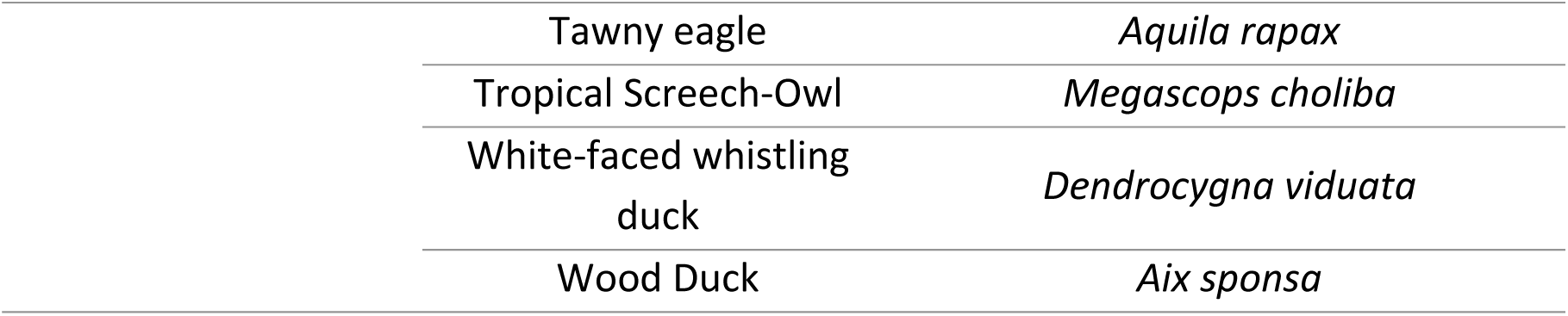
Classification of migration types among wild bird species affected by H5N1 in South America, 2022–May 2025.

## SUPPLEMENTAL MATERIAL

**Supplementary Figure 1.**
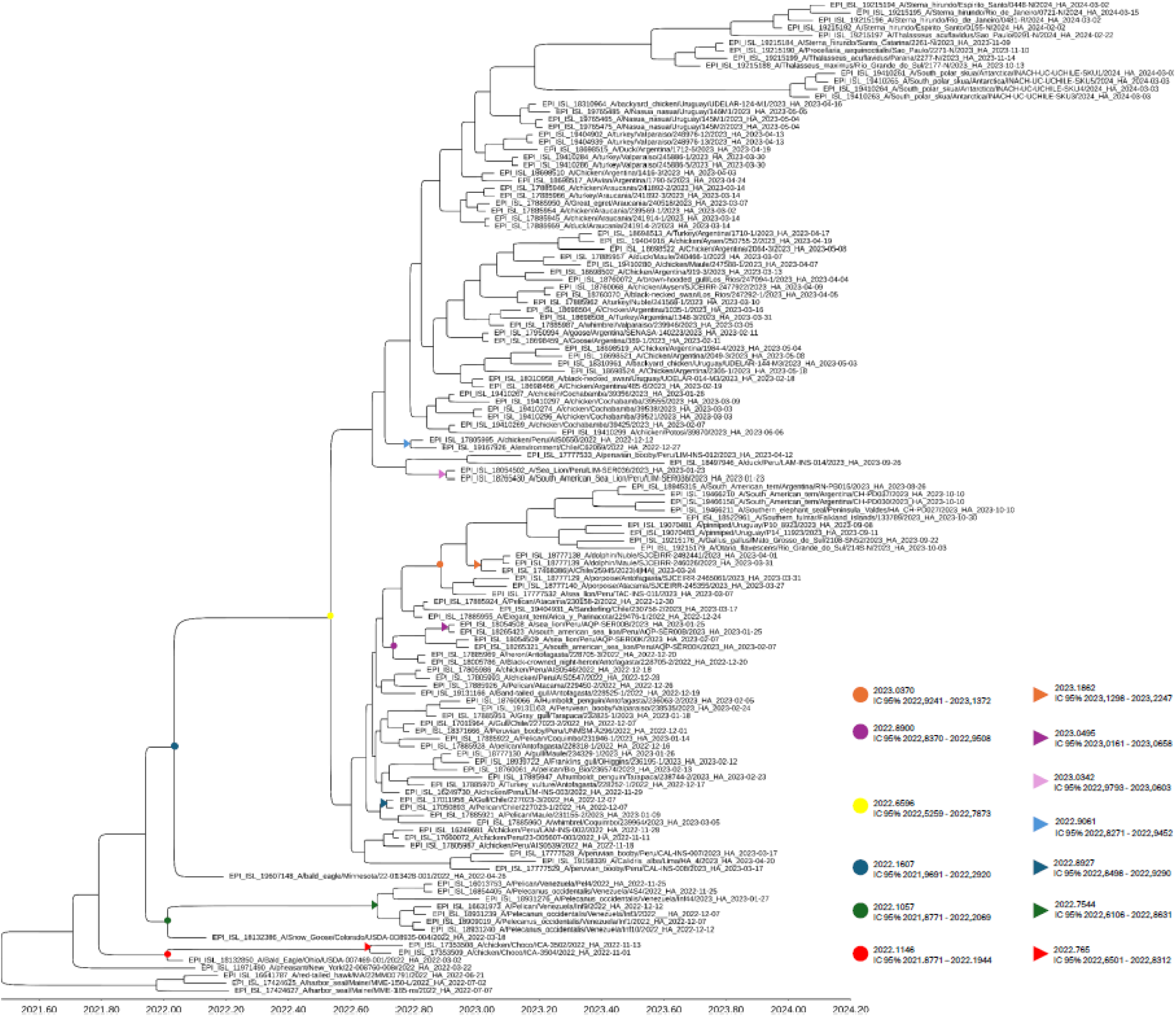
Time-calibrated phylogeny of Influenza A(H5N1) viruses based on the HA segment. The tree was inferred using BEAST v1.10.5.0 under a GTR substitution model, a relaxed lognormal molecular clock, and a Bayesian Skygrid coalescent prior. The analysis included 130 HA gene sequences sampled primarily in South America, with additional reference sequences from North America. Tip labels indicate host species, geographic origin, and sampling date. The x-axis represents time in decimal years, spanning from mid-2021 to 2024. Colored dots at internal nodes denote key epidemiological events: first introduction into South America (red), second introduction (green), first introduction into Chile (blue and light blue), first detection in marine mammals (Purple and light purple), and human infection (orange).

